# The interplay between proton diffusion across biological membranes and their biophysical properties highlights the role of defects in mixed lipid membranes

**DOI:** 10.1101/2024.02.06.579258

**Authors:** Ambili Ramanthrikkovil Variyam, Mateusz Rzycki, Anna Yucknovsky, Alexei A. Stuchebrukhov, Dominik Drabik, Nadav Amdursky

## Abstract

Proton circuits within biological membranes are at the heart of natural bioenergetic systems, whereas different biological membranes are characterized by different lipid compositions. In this study, we investigate how the composition of mixed lipid membranes influences the proton transfer (PT) properties of the membrane by following the excited-state PT (ESPT) process from a tethered probe to the membrane with time-scales and length-scales of PT that are relevant to bioenergetic systems. Two processes can happen during ESPT: the initial PT from the probe to the membrane at short timescales, followed by diffusion of dissociated protons around the probe on the membrane, and the possible geminate recombination with the probe at longer timescales. Here, we use membranes that are composed of mixtures of phosphatidylcholine (PC) and phosphatidic acid (PA). We show that the changes in the ESPT properties are not monotonous with the concentration of the lipid mixture; at low concentration of PA in PC, we find that the membrane is a poor proton acceptor. Molecular dynamics simulations indicate that at this certain lipid mixture, the membrane has the least defects (more structured and unflawed). Accordingly, we suggest that defects can be an important factor in facilitating PT. We further show that the composition of the membrane affects the geminate proton diffusion around the probe, whereas, on a time-scale of tens of nanoseconds, the dissociated proton is mostly lateral restricted to the membrane plane in PA membranes, while in PC, the diffusion is less restricted by the membrane.

## Introduction

Proton transfer (PT) in biological membranes is a key process in photosynthesis and aerobic respiration, which has been discussed ever since the advent of chemiosmotic theory in the mid-1960s and has been the subject of detailed studies in more recent years (1-5). It is recognized that PT in membranes is more complicated than in pure aqueous media (6, 7). Despite the efforts, the fundamental mechanism of PT across cellular biological membranes is still poorly understood due to the complex and multiscale nature of proton diffusion (PD) along distinct types of lipids and its interaction with the bulk phase (8-16). Heberle, Dencher, and co-workers discovered the ability of a membrane interface to act as a proton barrier, resulting in a delayed equilibration with the bulk phase and related long-distance (over µm) proton migration along the membranes (4, 17). However, the nature of the retaining barrier remains still poorly understood. Initially, it was believed that the slow rate of proton equilibration was caused by non-mobile pH buffers at the surface, such as lipids and amino acid groups. Later, it was argued that proton exchange between the surface and the bulk is retarded not simply by the immobile pH buffers but also by an interfacial potential barrier caused by electrostatic interactions (18, 19). Several models of the membrane-bulk interface have shown that both the pH-buffering capacity of the surface and the height of the interfacial potential barrier are responsible for determining the rate of proton exchange with the bulk and hence proton transport along the membrane; whether the excess proton can be mediated by the phosphate groups or to rapidly diffuse in the ‘shallow interface region’ (19-21). Among different lipids, it was shown that the anionic phosphatidic acid (PA) with a bare phosphate can have the ability to act as a proton collecting antenna from the bulk aqueous solution, i.e., to serve as the pH-buffering medium within the membrane; when the concentration of PA within the membrane was increased, the membrane was more susceptible to accepting a proton from the solution (22).

The capability of biological membranes to support long-distance lateral PD has been demonstrated for several types of membrane systems, starting from the early studies with the bacteriorhodopsin-containing membranes (1, 4, 5) until more recent studies showing a similar lateral PD coefficient by different membrane compositions (9, 23, 24). Interestingly, such studies included both membranes of low pH-buffering capacity: phosphatidylcholine (PC), phosphoethanolamine, or phosphoglycerol, as well as high pH-buffering ionic PA membrane. However, the molecular mechanisms of transport (both macro- and microscopic) remain unclear, in particular, the difference between ionic high pH-buffering PA and zwitterionic and low pH-buffering membranes remains surprisingly obscure. A difference that will be highlighted in this study.

In general, biological membranes are heterogeneous in terms of composition, which varies according to the type of cell (25, 26), and can significantly alter the (bio-)physical properties of the membrane (27) or create biologically significant patches known as domains (28). However, how the proton dynamics at the membrane interface will be affected by the mixing process of lipids is not well explored. Accordingly, our aim here is to investigate how a complex membrane composition can change its PT properties. Specifically, we wish to explore what is happening between the two mentioned extremes in terms of PT: the ionic and high pH-buffering PA membrane and the zwitterionic and low pH-buffering PC membrane. To do so, we used our developed photoacid probe, termed C12-HPTS, that can be tethered into the membrane and release a proton on the surface of the membrane assisted by hydrogen bonds between the probe and surface groups or water molecules (23). While time-resolved fluorescence provides information about the dynamics of a photo-dissociated proton on the membrane, our method can probe only relatively short timescales, up to 20-30 ns. Yet, on these timescales, protons can diffuse in aqueous media over distances up to 100-200 Å, which are typical between the respiratory enzymes on the membrane surface and thus of significant interest. We show that the composition of the membrane has a dramatic effect on PD around the probe following photo-dissociation, where we found anomalous proton dynamics at the membrane surface resulting in a poorer PT with small concentrations (≤33%) of PA within PC, followed by an increase in PT efficiency when more PA was added. To understand the membrane properties at low PA concentrations, we turned to molecular dynamics (MD) calculations. We found a similar anomaly as a function of the PC:PA ratio in the occurrence of defects. Our study is one of the first to correlate the membrane’s physical properties, i.e., packing and defects, with the proton dynamics on the membrane. The results shed new light on the complexity of proton dynamics in heterogeneous biological membrane environments.

## Results and Discussions

### 1. Steady-state measurements

Small unilamellar vesicles (SUV) of POPC (1-palmitoyl-2-oleoyl-glycero-3-PC) and POPA (1-palmitoyl-2-oleoyl-sn-glycero-3-phosphate) in different ratios along with the C12-HPTS probe were prepared (**Figure 1** for the chemical structure of the lipids and the probe along with the ratios used in this study). As shown in **Figure 1**, whereas the long alkyl tail part is identical for both lipids, POPA possesses an anionic bare phosphate group while POPC has a zwitterionic head group. The PO lipid system was chosen due to its low melting point, thus ensuring that the formed vesicles are in their liquid state. The formation of a mono-dispersed solution of liposomes was confirmed using dynamic light scattering (DLS) with a uniform size for all the different vesicles prepared (**Figure S1**). We further used confocal fluorescence microscopy of the membranes to confirm the uniform incorporation of the C12-HPTS probe into the membranes (**Figure S2**).

**Figure 1.**
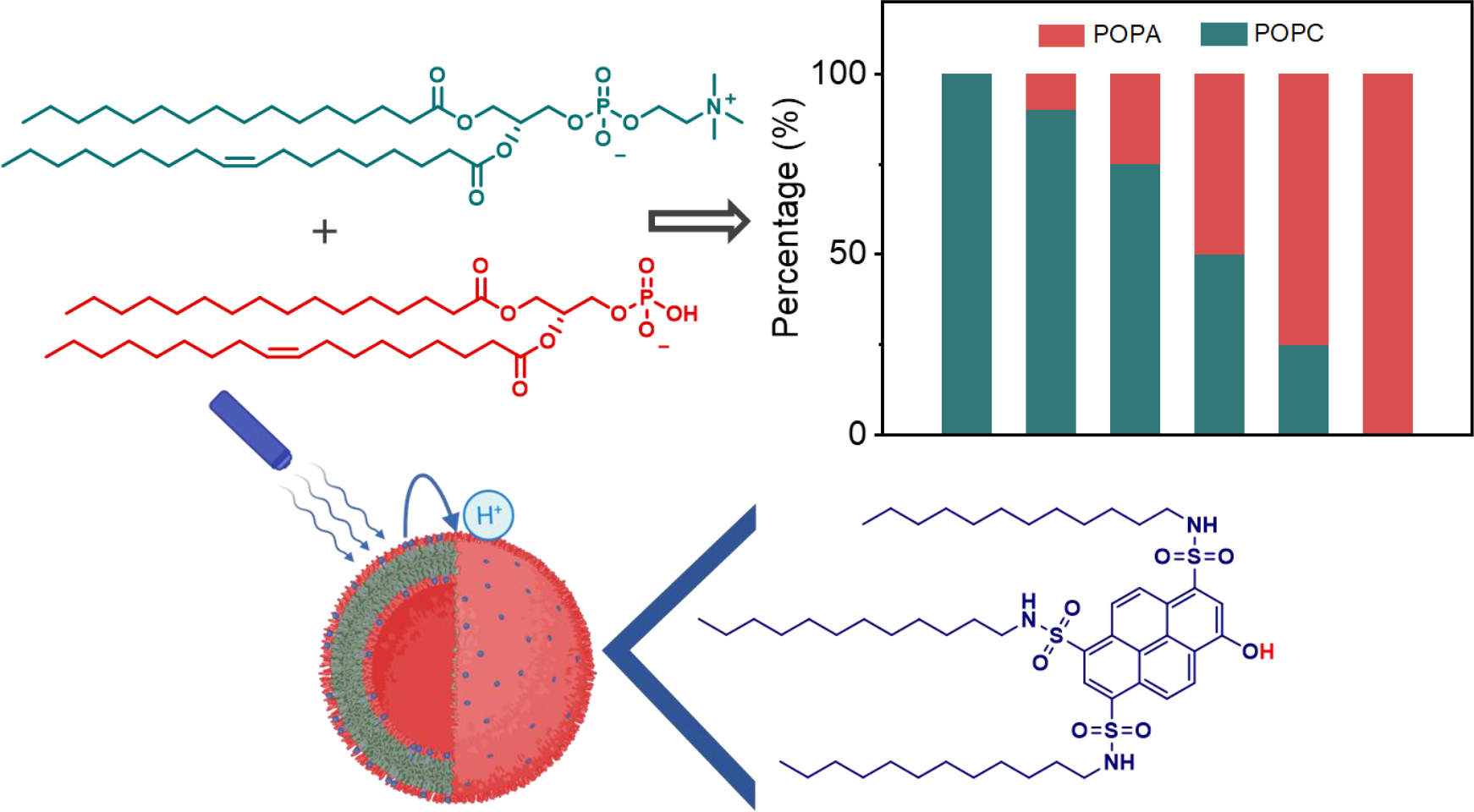
Top panel: Molecular schematic of POPA and POPC lipids and the ratios used for making the SUV’s. Bottom panel: Molecular schematic of C_12_-HPTS and its incorporation into an SUV together with a schematic of a light-trigger PT event from the photoacid to the membrane.

As for the optical properties of the C12-HPTS probe within the membrane, and in line with any Brönsted photoacid, the optical absorbance of its protonated (ROH) and deprotonated (RO^-^) forms are at different wavelengths. To use the photoacid probe for investigating the PT processes on the membrane surface by using the excited-state proton transfer (ESPT), it must be excited in its ROH form (29):

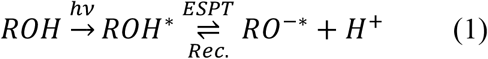

The UV-Vis absorption of C_12_-HPTS in all of the membranes investigated here shows a predominant ROH peak at pH≤7.4 (**Table S1** and **Figure S3**). The UV-Vis absorption can be used to evaluate the p*K*_*a*_ of the probe using pH titration (**Figure S3**). We showed previously that the p*K*_*a*_ of the photoacid probe within POPC membranes is ∼8, whereas the p*K*_*a*_ within POPA membranes is higher, at ∼10, and that the peak positions are undergoing some blue shift at anionic membranes (23). In line with these results, we found here that the p*K*_*a*_ of C_12_-HPTS in all of the mixed PC:PA membranes is between these values, and the more negatively the membrane charge (**Figure S4** for the zeta-potential of the vesicles at different ratios), the higher the calculated p*K*_*a*_ (**Table S1**). It should be noted that we can only measure the apparent p*K*_*a*_ values based on the pH in the solution and not the exact proton concentration on the surface of the membrane.

In the excited state ROH*, the p*K*_*a*_ drops to low values (30) and ROH* quickly (<1ns) deprotonates. The formation of RO^-*^ state is evident in the fluorescence spectra, where a predominant peak corresponding to RO^-*^ along with that of ROH* is observed (**Figure 2a**). In line with the optical absorbance measurements, the more negatively charged the membrane is, the more blue-shifted the position of the fluorescence peak (**Table S1**). The ratio RO^-*^/ROH* (**Figure 2b**) is an indicator of the ESPT processes (30). Surprisingly, as a function of membrane composition, the ratio RO^-*^/ROH* is not a simple monotonous function, as Figure 2b shows, but has an anomalous minimum at low %POPA concentrations, which already indicates a non-trivial nature of the PT process at the membrane surface.

**Figure 2.**
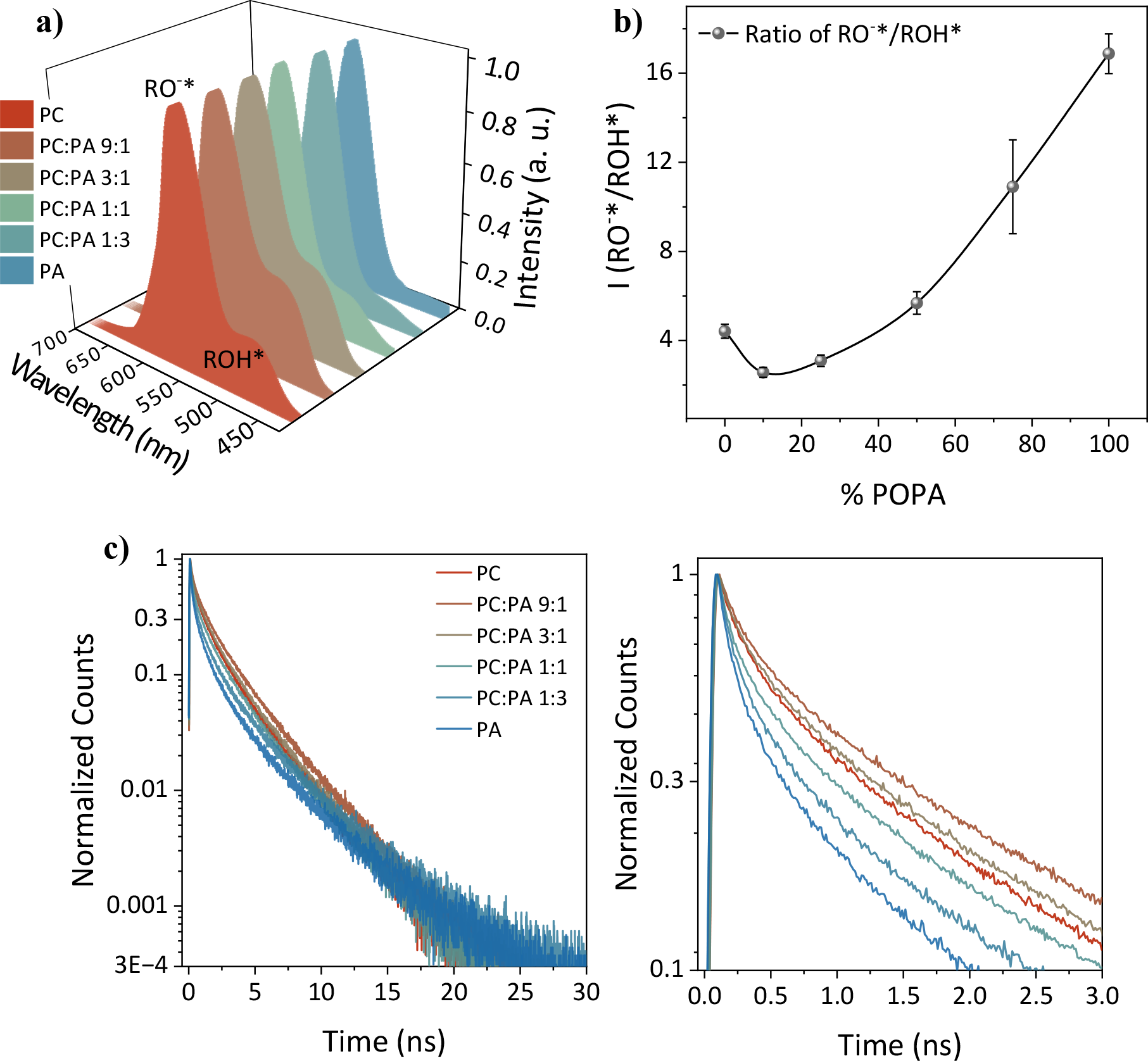
a) Steady-state along with b) the extracted RO^-*^/ROH* ratio, and c) time-resolved fluorescence measurements of C_12_-HPTS within SUV’s at various lipid compositions taken at pH 7.4, λ_ex_=410 nm, λ_em_=470 nm (for panel c). The right panel in (c) shows a zoom-in of the first 3 nanoseconds in the left panel.

Upon constant illumination in a steady-state measurement, the intensity of RO^-*^ peak is defined by the stationary state population of RO^-*^; the latter is defined by the balance of formation and deactivation processes, so that the ratio RO^-*^/ROH* can be described as:

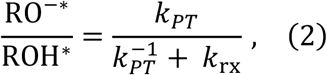

where *k*_*pT*_ is the rate of deprotonation of ROH* state (rate of formation of RO^-*^, marked as *ESPT* in Eq. (1)), 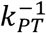is the reverse protonation rate, i.e., recombination (*Rec*. in Eq. (1)), due to geminate protons and the protons of the medium, and *k*_rx_ is the RO^-*^ deactivation rate due to both radiative and non-radiative processes resulting in the ground state RO^-^. Accordingly, a change in the membrane composition might influence RO^-*^/ROH* in various ways. However, in our steady-state measurements, we cannot extract the parameters of the ESPT process and the dynamics of the geminate protons around the C_12_-HPTS probe attached to the membrane. This can be achieved however in time-resolved experiments.

### 2. Time-resolved measurements

To further understand the processes occurring in the excited-state of the C_12_-HPTS probe within the membrane, we turned to time-resolved fluorescence measurements. As shown in Eq.(1) and discussed above, two processes are happening during the excited-state that influence the transient populations of [ROH*] and [RO^-*^]: The ESPT, with membrane acting as the proton acceptor, that happens immediately after excitation, and the subsequent geminate recombination process that is prominent at longer (>1 ns) timescales. The geminate recombination process reflects the dynamics of the dissociated proton around the probe in the membrane (28) and is of our particular interest.

The short timescale fluorescence decay of ROH* is mainly due to deprotonation; thus, it reflects the rate *k*_*pT*_ of the ESPT process. As seen in **Figure 2c** and from the extracted values for a three-exponent fitting of the decays (**Table S2**), the short-time decay rates change with the composition of the membrane in a similar non-monotonous way as in the steady-state experiment.

To quantitatively relate the short- and long-timescale decays in **Figure 2c** to the ESPT processes, we use our model of geminate recombination (31). The model predicts that at short times, the decay of the ROH* state is linear in time, resembling the initial stage of an exponential decay process, and the relation to *k*_*pT*_ is as follows:

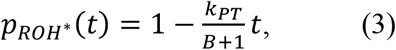

where *k*_*pT*_ is the dissociation constant of the photoacid, and B is a Boltzmann factor reflecting the effect of the negative charge of the dissociated photoacid on the geminate proton dynamics (see further discussion in Ref. (31)).

At long timescales, the fluorescence decays according to a power law:

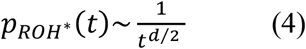

where *d* is the dimensionality of the medium around the probe in which the geminate proton diffusion occurs. Accordingly, in a log-log plot shown in Figure 3, the long-time decay appears as linear, with a slope of *d*/2. Thus, at the long timescale, the fluorescence decay reflects how the dissociated proton migrates around the probe before recombination takes place. Using the model, the three independent parameters, *k*_*pT*_, B, and *d* can be determined using the short and long timescale asymptotics of the fluorescence decay. The short timescale rate *k*_*pT*_, and the long timescale *d* parameter are two key parameters that characterize ESPT: *k*_*pT*_ describes how fast the injection (dissociation) of the proton into the membrane surface, and the dimension *d* describes how the dissociated proton moves around the probe on the membrane surface.

**Figure 3.**
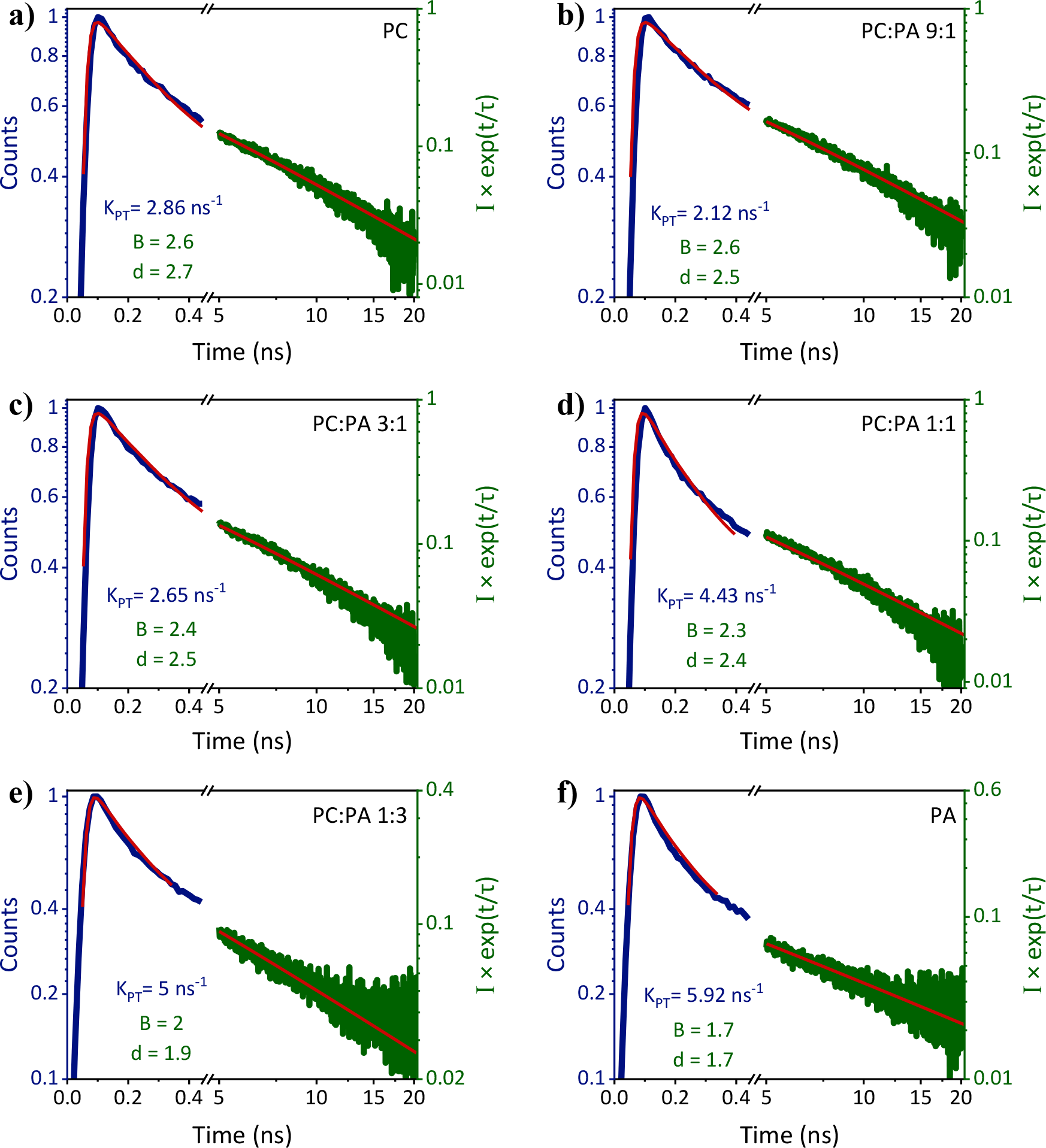
Fitting of the time-resolved fluorescence measurements at short timescales (blue) and long timescales (green) of: a) PC, b) PC:PA 9:1, c) PC:PA 3:1, d) PC:PA 1:1, e) PC:PA 1:3, and f) PA.

Here, we aim to understand how *k*_*pT*_ and *d* depend on the membrane composition. Accordingly, we have fitted our experimental results according to our model at the two described time regimes (**Figure 3**), and extracted *k*_*pT*_ at the short timescales and *d* at longer timescales (text in **Figure 3** and **Table S3**).

On the surface of POPA, the extracted dissociation rate *k*_*pT*_ is fast (5.92 ns^-1^), and the dimensionality parameter *d* is close to 2 (d=1.7), which indicates that on the timescale of 10-20 ns, the dissociated proton is apparently restricted to the membrane and moves along the membrane surface. POPC showed a slower *kPT* (2.86 ns^-1^) and *d* values closer to 3 (d = 2.7), which means the dissociated proton is less restricted to the membrane surface on the 10-20 ns timescale (it is important to note here that while the time-resolved decay of C_12_-HPTS in pure POPC and POPA membranes is similar to what was reported in Ref. (23), the use of different models resulted in different extracted values). Unrestricted PD on timescales of ∼20 ns corresponds to distances of more than 100 Å. For POPC, the released proton is not exclusively restricted to the membrane surface, though it does not exclude a barrier on the larger distance scale from the membrane surface. For POPA, the proton is more restricted to the surface of the membrane. This simplified picture is probably more complicated, and there might be some restrictions to the surface in POPC, and the 2D diffusion on the POPA surface might have some underlying structure that can be imagined as a network of molecular water wires between the phosphate groups. Our conclusion here is supported by computational studies emphasizing the role of the phosphate groups in mediating the excess proton (21).

After defining the extremes in our system, i.e., how we can describe the PT process on the surface of pure POPC and POPA using our time-resolved measurements and the use of the described model, we reach now to what is happening in the mixed membrane heterostructures.

### 3. The ESPT process at different membrane compositions

In this chapter, we will discuss several different parameters that can influence PT and PD, and how they can influence the measured observables in a monotoneous or non-monotonous way.

*The role of k*_*pT*_: The straightforward parameter that is being influenced is *k*_*pT*_, which we already know is different for PC membranes vs. PA membranes (23). Qualitatively, one expects the increase of *k*_*pT*_ for POPA membrane, as more proton accepting groups are present, and the ratio RO^-*^/ROH* and the extracted *k*_*pT*_ are expected to increase. However, both the steady-state RO^-*^/ROH* and the *k*_*pT*_ from the time-resolved measurements are exhibiting a similar non-monotonous trend i.e., a reduction in RO^-*^/ROH* and the *k*_*pT*_ upon adding a small amount of PA into PC, followed by an increase upon adding more PA, until reaching the high RO^-*^/ROH* and *k*_*pT*_ values of POPA (**Figure 2b** and **Figure 4**). Hence, it is safe to conclude that the observed anomalous non-monotonous changes are mainly due to a change in the short timescales of the process, i.e. ESPT from the photoacid to the membrane. It is important to mention that the magnitude of change is different, whereas the magnitude in the change of RO^-*^/ROH* as observed in the steady-state measurements (**Figure 2b**) is somewhat larger than the change in the extracted *k*_*pT*_ values (**Figure 4**). Hence, other processes are also influencing the RO^-*^/ROH* ratio observed in the steady-state measurements.

**Figure 4.**
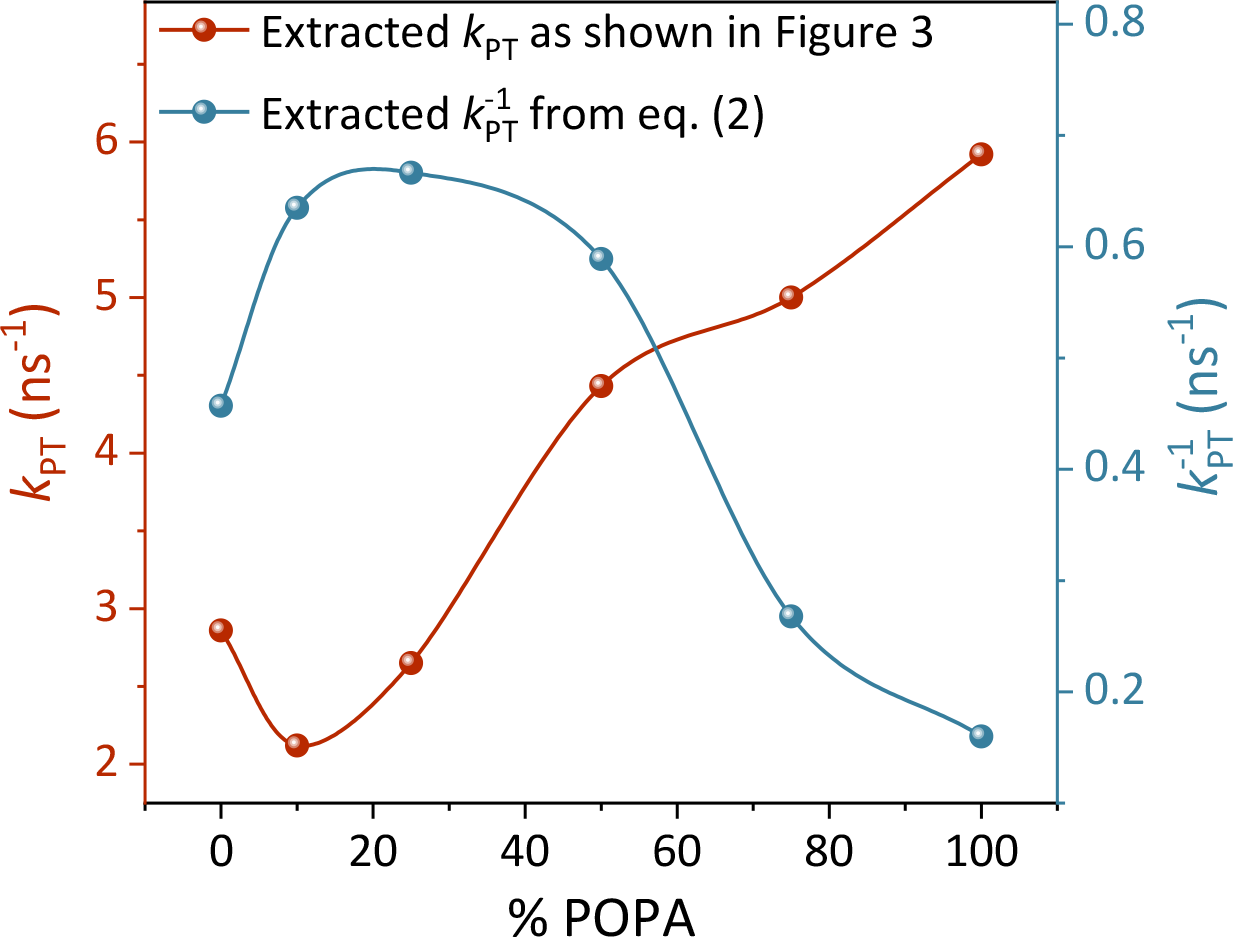
The extracted *k*_*pT*_ values from the time-resolved measurements using the model described in Figure 3 and the extracted 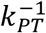 from Eq. (2).

*The role of k*_*rx*_: A change in the membrane composition might well influence *k*_rx_; the combined radiative and nonradiative deactivation of RO^-*^ (Eq. (2)), and in turn will result in the observed non-monotonous change of RO^-*^/ROH*. To probe it, we measured the decay of the RO^-*^ species, showing identical decay curves with a *k*_rx_ of 0.2 ns^-1^ for all membranes used in this study (**Figure S5**). Hence, we should exclude this change as the main cause for the observed non-monotonous change of RO^-*^/ROH*. Furthermore, considering the slow decay of RO^-*^, *k*_*pT*_ is significantly (up to 10x) higher than *k*_rx_.

*The role of* 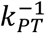: The additional parameter that is important for the observed non-monotonously change in RO^-*^/ROH* and its magnitude is 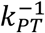, i.e., the protonation of RO^-*^ as a function of %PA in the membrane. In the time-resolved measurements, it manifests in the magnitude of the long-lived tail at long time scales (a3 in **Table S2**). As shown in the table, this component also shows a non-monotonous change as a function of %PA in the membrane. Using Eq. (2) and the extracted *k*_*pT*_ and calculated *k*_rx_, we can estimate 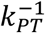 (**Figure 4**). The figure shows a non-monotonous behavior where 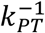 rises at low concentration of PA within PC followed by a decrease at high PA concentrations. As discussed, 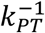 is complex and dependent on both protons that are coming from the medium and geminate protons from the membrane:

1) For protons coming from the medium, we expect a monotonous change due to the charge of the membrane that becomes more negative with the increase in %PA, resulting in an increase in 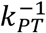 as a function of %PA in the membrane (unlike **Figure 4**). To directly estimate the role of protons coming from the medium, we complemented our measurements at neutral pH (**Figure 2a**) with steady-state measurements at lower pH values (**Figure S6**). As expected, RO^-*^/ROH* is decreasing as a function of lowering the pH. However, this decrease is very mild considering the orders of magnitude change in proton concentration of the medium, thus highlighting the relatively minor role of this parameter. Moreover, and importantly, the non-monotonous change of RO^-*^/ROH* is present at all pH values.

2) For geminate protons coming from the surface of the membrane, the situation is more complicated. According to the described mechanism, following excitation, ROH* deprotonates, and the geminate proton can be accepted by the membrane and diffuse away. This PD capability of the membrane will highly influence the geminate proton recombination that is dependent on their concentration around the probe. Accordingly, it is safe to assume that the membrane composition will influence this parameter. Indeed, for POPA we show a fast (lateral) PD of the geminate proton from the probe, thus resulting in a low 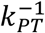, whereas for POPC 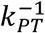 is higher due to the restricted motion (the extremes in **Figure 4**). The geminate recombination of protons can be influenced by the discussed dimensionality of PD. However, this dimensionality is monotonous as a function of %PA (**Figure 3** and **Table S3**). Hence, it is safe to claim that the anomaly in the membrane properties that cause the non-monotonous behavior of *k*_*pT*_ is also affecting the capability of the membrane to reprotonate RO^-*^.

The main question remaining now is which type of anomaly in the membrane properties can result in the observed changes of *k*_*pT*_ and 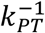, which leads us to the next section.

### 4. MD simulations

In this section, we will discuss what makes the membrane at PC:PA ratios of <3:1 a bad proton acceptor for protons that are being released on its surface and at the same time more efficient for geminate recombination? Indeed, one would expect a monotonous increase of *k*_*pT*_ with the increase in the number of proton-accepting phosphate groups. To address this question, we turned to MD simulations of membranes at different PC:PA ratios. At first, we used both coarse-grain and all-atom simulations to examine the interaction between PC and PA lipids and to probe whether substantial domain formation could be observed, where we found no indication for domain formation (**Figures S7-8**). Accordingly, in our calculations, we focused on several membrane-associated parameters at different PC:PA ratios (**Figure 5**) as follows (see also section ‘Molecular Dynamics membrane systems validation’ and **Figure S9** in the Supporting Information for a benchmark analysis of the obtained value with previous studies):

**Figure 5.**
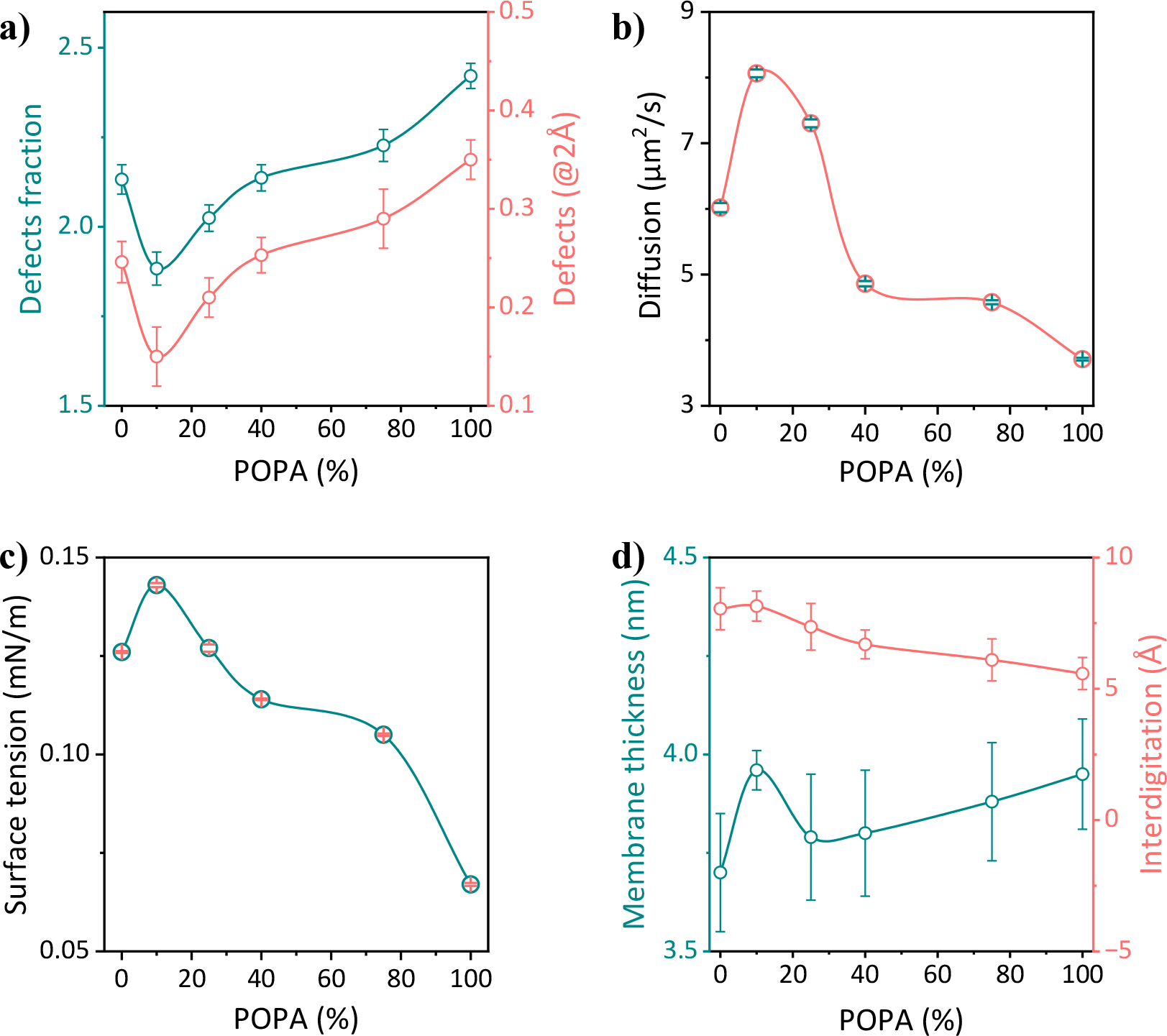
MD simulation of various membrane properties as a function of lipid composition (% of POPA in POPC): a) Defects – showing two parameters, the first is the ratio of exposed acyl chain regions to polar regions (cyan curve and left y-axis) and the other is representing the ratio of regions to which a sphere of radius 2 Å could be fitted (red curve and right x-axis); b) Lateral diffusion of lipids within the membrane; c) Surface tension of the membrane; d) The membrane thickness (cyan curve and left y-axis) and interdigitation of the acyl chains (red curve and right x-axis).

1) Defects (**Figure 5a**): We will start with the membrane parameter that shows a perfect correlation to our experimental result, which is defects quantity within the membrane structure. To quantitatively approach defects, we used a two-level approach: the first was to calculate the ratio of exposed acyl chain regions to polar regions, and the second was to calculate the level of accessibility of exposed acyl chain regions by a sphere with a radius of 2Å (see snapshots in **Figure S10**). The parameters obtained from both levels resulted in a similar outcome; adding PA into PC at 9:1 of PC:PA induces a major change in the membrane making it less defective, i.e., more structured and unflawed. This is evident by the lower exposed acyl chain to polar regions, thus suggesting a more condensed structure, as well as in the less accessibility with the 0.2 nm sphere.

2) Diffusion of lipid molecules (**Figure 5b**): This parameter also shows a similar trend with the experimental result, and it is probably derived from the discussed defects in the previous section. While this work targets PT on the surface of the membrane and PD, we should also consider the diffusion of the lipid molecules themselves. As shown in **Figure 5b**, the diffusion of lipids within the membrane containing the PC:PA ratios of 9:1 and 3:1 is faster than all other membranes used in this study. As discussed above, in these exact ratios, the membrane is the least defective. These two results agree with each other, whereas a more structured and unflawed membrane allows faster lipid diffusion. The question of why a more structured membrane having a faster lipid diffusion results in having a poor PT will be tackled below.

3) Surface tension (**Figure 5c**): This parameter is also related to the first one. It was found that surface tension is related to how the membrane is structured and to the abundance of defects, whereas some defects can result in lower surface tension (32). Also in our calculations, the most structured and unflawed membrane at the PC:PA ratio of 9:1 is also the membrane with the largest surface tension, and as can be observed in the figure, the overall trend matches the ones in **Figures 5a** and **4b**.

4) Membrane thickness and interdigitation (**Figure 5d**): Unlike the previously discussed parameters, membrane thickness and interdigitation show a more linear trend as a function of the PC:PA ratio. Higher interdigitation results in a thinner membrane thickness, as for POPC, and vice versa for POPA. Also here, we have some discrepancy specifically for the 9:1 PC:PA ratio that shows somewhat higher thickness but with similar interdigitation as for POPC. While this discrepancy might be explained by domain formation, we did not find a strong indication for domain formation neither in our MD calculations (**Figures S7-S8** and text within) nor in confocal fluorescence microscopy measurements of vesicles containing low %PA (**Figure S11**).

Summarizing, it is well evident that the addition of a small amount of PA into PC in a 9:1 ratio (PC:PA) considerably changes a set of parameters related to the structure of the membrane. At this specific ratio, the membrane is most packed (resulting also in the lowest area per lipid, **Figure S12**) and structured, having the highest surface tension, and the least defective among all other lipid compositions. The unique structural properties of this membrane are probably due to attractive forces between the PA and PC lipids (34, 35). Nevertheless, our experimental observations that this specific membrane is also a poorer proton acceptor, exhibiting the least efficient ESPT from the photoacid probe, might appear counterintuitive. Below we discuss the different membrane parameters associated with its capability to transport a proton that helps to rationalize the result.

### 5. Why is a well-structured membrane a poor proton acceptor?

To answer this question, we first need to understand the various PT processes that can occur on the surface of a membrane and their interactions with the bulk medium surrounding the membrane. POPC and POPA membranes are quite different. For POPA membranes, with their exposed phosphate groups to the aqueous solution, the surface of the membrane can serve as a good proton acceptor for both protons that are released on the surface of the membrane from the tethered photoacid (23), and for protons coming from the solution, i.e., the ‘proton antenna effect’ (22, 36). As discussed above, protons accepted by POPA can be retained and diffuse on the surface of this membrane for timescales of 10-20 ns (corresponding to distances of >100 Å). On POPC membranes, we find the probe releases the proton less efficiently than on the POPA surface. Naturally, the zwitterionic POPC surface is a less efficient proton acceptor than the negatively charged POPA surface. But still, POPC is an efficient proton acceptor. While we expected that the gradual addition of POPA into POPC would make the membrane progressively a better proton acceptor, we found instead an anomalous, non-monotonic behavior at low PA:PC ratios. Surprisingly, the addition of a small amount of PA to PC makes the membrane an even poorer proton acceptor. As MD simulations indicate, this surprising non-monotonous behavior correlates with the change in the structural properties of the mixed membrane, described here in terms of defects. With the addition of a small amount of PA, the less defective, more structured membrane becomes a poorer proton acceptor, i.e. having lower ESPT rate *k*_*pT*_, despite the added proton accepting phosphate groups in the membrane. Additionally, we also observe that the poorer *k*_*pT*_ is accompanied by a higher 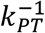, meaning the poor ability of the membrane to support PD away from the probe is resulting in a more efficient proton geminate recombination.

This is surprising, and we can only speculate the ‘*why*’ question. One reason might be related to the discussed dimensionality and the suggested PT mechanism that is assisted by water wires on the surface of membranes. As shown in **Figure 5a**, the membrane that exhibits the most defects is POPA. Accordingly, we can claim here that such defects are important for the efficient retention of protons on the surface of POPA and its lateral PD. Adding PA into PC helped organize the membrane to a less defective state, but as can be observed in **Figure 3** and **Table S3**, it did not result in a major change in dimensionality. From this observation, we can conclude that having PA lipids in the membrane does not necessarily assist in PT to the surface of the membrane, but they can even suppress the PT capabilities of the membrane by interfering with any possible water wires on the surface of PC lipids. Increasing the PA concentration shifts the balance toward the ‘PA effect’ – a more defective membrane, change in dimensionality, and fast PT from the photoacid to the membrane. To have some insight into the presence of water molecules with respect to the presence of defects in the membrane, we turned back to our MD simulations. In the simulations, we found that on average, areas with high concentrations of water molecules can be correlated to some extent with the position of defects (**Figure S13**), which might explain why a ‘defective’ membrane is a better proton mediator. It might be also that the less defective membrane changes the position of the C_12_-HPTS probe on the surface of the membrane which can also induce a change in the ESPT process.

Our findings here are also in line with our previous study on the role of adding surfactants on the PT of biological membranes (37). In our previous study, we observed an improvement in the measured PT rate as a function of adding surfactants, whereas the larger the surfactant concentration the better the PT. While previously we speculated it is the major damage of the surfactant to the membrane structure that results in enhanced PT properties, i.e., the escape of protons from the surface to the water, we now understand that defects also had a role in promoting PT in that study.

Lastly, our new results here together with the calculations are pointing to an interesting decoupling between lipid diffusion to PD, thus highlighting the different mechanisms for each type of diffusion. In lipid diffusion, a well-structured membrane with the least defects allows the rapid diffusion of phospholipids in it. On the other side, PD is dictated by a different mechanism, usually described by the Grotthuss mechanism of having a percolating ‘wire’/network of hydrogen bonds that assist in promoting PD through numerous proton hopping steps. For membranes, such a network can be composed of moieties within the phospholipid, primarily the phosphate group in our case, as well as water molecules that are present on the surface of the membrane. Our finding here that more defects within the membrane resulting in an improved PT suggests that the percolating network on the surface needed for a Grotthuss-type PD process is better, and the likely cause for it is that defects can allow more water molecules to be incorporated into the membrane structure. The correlation between the number of water molecules on a biological surface to its ability to diffuse protons in a Grotthuss-type PD process is well-studied (38-45).

## Conclusions

PT on the surface of biological membranes is a complex phenomenon. To investigate it, we used several membranes with different ratio of PC:PA and monitored the ESPT process from a tethered C_12_-HPTS photoacid. Using steady-state and time-resolved fluorescence measurements, we found an anomalous behavior in the ESPT process, showing that the composition of the membrane has a dramatic effect on PD around the probe following photo-dissociation. On a timescale of tens of nanoseconds, the dissociated proton moves mostly laterally on the surface of PA membranes, whereas in PC membranes, the diffusion is less restricted by the membrane. The anomaly is observed at low concentrations of PA within PC, manifested in a non-monotonous change in the experimental results going from pure POPC to pure POPA. We concluded that the observed anomaly is due to both the initial ESPT between the probe and the membrane and the subsequent geminate recombination process. To understand the reason behind the peculiar finding, we turned to MD simulations, where we found that the level of defects within the membranes at different PC:PA ratios match perfectly the experimental results concerning the PT rate constants. In our discussion, we tried to rationalize why a less defective, i.e., more structured and unflawed, membrane would exhibit the poorest PT properties. Interestingly, the results suggest that a well-structured membrane inhibits PT while a more defective one promotes the PT process. The timescales of PT/PD probed here correspond to proton diffusion over long distances exceeding 100 Å, which are typical distances between the enzymes of the respiratory chain. Thus, our results provide new insights into possible mechanisms of proton transport in bioenergetic circuits.

## Materials and Methods

The detailed Materials and Methods section is in the Supporting Information section.

### Experimental Session

SUVs of different lipid membranes were prepared for experiments using the extrusion methodology through a polycarbonate membrane of 200 nm-sized pores. The SUVs were used for all the spectroscopic measurements including steady-state and time-resolved fluorescence experiments using an ultrafast laser system. All the measurements were carried out at room temperature (∼23°C).

### MD Investigations

Membrane models were created using CHARMM-GUI membrane builder(46). The systems contained different lipid ratios and each consisted of 400 lipids neutralized with Na^+^ ions to balance the net charge. The MD simulations were performed using GROMACS (version 2021) software (47) and CHARMM36 force field(48). Prior to production runs, the membrane systems underwent energy minimization and equilibration using the standard CHARMM protocol (46) to achieve stability and reproducibility in the simulations. The area per lipid (APL) and membrane thickness (MT) were determined using a custom MATLAB script. Bending rigidity was calculated using the real space fluctuation (RSF) method(49), while the lateral diffusion coefficient was determined through Einstein’s relation(50). Interdigitation was evaluated using MEMBPLUGIN(51), and surface tension was calculated using the Irving-Kirkwood method(52). The defects in the bilayers were analyzed using a protocol based on the accessibility of the acyl chains(53), including the calculation of occupancies and the partitioning of the simulation box into grids.

## Acknowledgments

N.A. and A.A.S thank the United States - Israel Binational Science Foundation (grant number: 2018239 (NA) and A19-3374 (AAS)) for financial support. N.A. thanks the Lower Saxony – Israel Research Cooperation (grant number ZN3625) for financial support. M.R. thanks the National Science Centre, Poland (grant number 2022/45/N/NZ9/02130) for financial support and Wroclaw Centre for Networking and Supercomputing (http://www.wcss.pl) for providing computing resources (grant number 274).

