## Supporting information for "The interplay between proton diffusion across biological membranes and their biophysical properties highlights the role of defects in mixed lipid membranes"

### Materials and Methods

**MD Investigations:** To investigate the behavior and properties of POPA/POPC bilayers, several membrane systems were created using the CHARMM-GUI membrane builder, consisting of POPC and POPA mixtures in the following ratios: 1:0, 0.9:0.1, 0.75:0.25, 0.6:0.4, 0.25:0.75, 0:1 (1). Further, the lipids were solvated with the TIP3P water model, including 58 water molecules per lipid to ensure sufficient lipid solvation (2). All PA/PC systems contained 400 lipids and were neutralized with $Na^+$ ions to balance the total net charge. The GROMACS (version 2021) software (3) and CHARMM36 force field were employed to perform MD simulations (4).

First, the membrane systems were minimized using a steepest decent algorithm for energy minimization. Further, the NVT (constant Number of particles, Volume, and Temperature) ensemble was applied according to CHARMM protocol with the leap-frog integrator and a 1 fs timestep. Under the NPT (constant particle Number, Pressure, and Temperature) dynamics a 2-fs time step was enabled. Simulations were carried out using a Nose-Hoover thermostat at T = 303.15 K with a time constant $\tau t$ = 2 ps (5, 6). A semi-isotropic Parrinello-Rahman barostat with p = 1 bar and $\tau p$ = 5 ps was employed for pressure coupling (7). Under the equilibration steps, positional and dihedral restraint potentials were utilized, and their force constants were gradually decreased. Bond distances were maintained using the LINCS algorithm (8). In the production run the hydrogen mass repartitioning technique enabled 4 fs time step integration (9). The van der Waals interactions were gradually switched off over 1.0 - 1.2 nm using a force-based switching function (10), while long-range electrostatic forces were evaluated by the particle mesh Ewald (PME) method with a 0.1 nm grid spacing and 1.2 nm cutoff (11).

Area per Lipid and Membrane Thickness: Both membrane area per lipid (APL) and membrane thickness (MT) were determined using a custom MATLAB script. The point position for each lipid molecule was calculated as the midpoint between P and C2 atoms' position. Those points were followed by Voronoi tessellation to obtain the individual APL for each lipid molecule in every simulation time step. Obtained APL values were histogrammed and the average value was determined by fitting the Gaussian function. MTP-P was calculated as the difference between average height (z-axis) values of phosphorus in opposite leaflets.

Bending Rigidity: Real-space fluctuation (RSF) method was used to determine the bending rigidity of investigated systems (12). Specifically, a probability distribution for splay is determined for all lipids over all analyzed time steps. Lipid splay is defined as the divergence of the angle formed by the directors of neighboring lipids providing that they are weakly correlated.

Lateral Diffusion Coefficient: The Diffusion Coefficient Tool plugin was used for the determination of lipid molecules' lateral (2D) diffusion (13). It is calculated using Einstein's relation with the mean square displacement (MSD) of the chosen molecular species. Therefore, the time-dependent diffusion coefficient D(t) can be calculated via the given relation:

$$D(t)=\langle|r(t)-r(0)|^2/2dt\rangle$$

Where r(0), r(t), denote the particle position at arbitrary time origin and at time t, respectively. The lateral diffusion of given lipid species was calculated in xy dimensions (d) based on phosphorous atoms' positions. Taking into account common systematic errors in diffusion coefficient determination, trajectories were thoroughly analyzed in 50 ns sets as Bulow et al. suggested in their work (14).

Interdigitation: The interdigitation parameter was determined using MEMBPLUGIN (15). The obtained parameter is defined as the width of the region of mass overlap, reflecting the interdigitation between the opposite leaflets in the system.

Surface Tension: The GROMACS software's built-in function *gmx energy* was utilized to calculate the surface tension of the PA/PC membranes. Membrane surface tension was computed with a pressure tensor ($P_{xx}$, $P_{yy}$, $P_{zz}$) based on the Irving-Kirkwood method (16).

$$\gamma\approx L/2\ \langle P_{zz}- (P_{xx}+P_{yy})/2\rangle$$

Here L denotes the length of the simulation box in the z dimension, while the brackets represent an ensemble average given from gmx energy.

Defects analysis: A protocol implemented by Boyd et al. was used for defect determination by acyl chain accessibility (17). Briefly, the occupancy of each bilayer atom (from lowest to highest) is calculated based on the Van der Waals radius. For headgroup atoms is marked as polar, otherwise as a tail. The headgroup is defined as all atoms from top to down including the C2 atom. This is followed by dividing the simulation box into square grids with a grid spacing of 0.5 Å. For each grid position, a line is led and id marked as head or tail depending on the first collision with atoms. As a result, both a 2D map of acyl chain accessibility and a fraction of head-to-tail grid points are obtained. This is followed by sphere probe analysis in order to ignore small gaps in headgroup coverage. For each point, a sphere probe of radius ranging from 0.1 nm up to 0.5 nm is set. If all the grid points in the range of this probe are classified as tails, those grid points are assigned as defects. In this way, only hydrophobic patches with given sizes and shapes are classified as true defects, which has the advantage over a previously determined 2D map of acyl chain accessibility.

**Preparation of small unilamellar vesicles (SUVs):** All the lipids were purchased from Avanti Polar Lipids and used without further purification. The probe was synthesized according to the procedure in the previous paper (18). Lipids (2 mM) were mixed with the probe in chloroform at a ratio of 100:1. The solvent was removed using a rotary evaporator and dried further under vacuum overnight. The formed dried lipid film was rehydrated and dissolved in 5 mM phosphate buffer (2 ml). Then the resulting solution was extruded 31 times through a polycarbonate membrane of 200 nm-sized pores. The extruded solution was stored at 4°C and the pH of the solution was adjusted using NaOH or HCl.

**Steady-state spectroscopic measurements:** Steady-state UV-visible absorption experiments were recorded by Agilent Cary 60 and the steady-state fluorescence measurements were performed using an FS5 spectrofluorometer (Edinburg Instrument). DLS and zeta potential measurements were carried out on Malvern Instruments Zetasizer Nano ZS. The refractive index and absorption of liposomes are assumed to be 1.450 and 0.010 respectively. The temperature of measurement was 25°C and the measurement was carried out using ZEN0040 cuvette.

**Time-resolved fluorescence measurements:** Time-correlated single photon counting experiments were carried out using a CHIMERA spectrometer (Light Conversion) with an excitation at 420 nm for $C_{12}$-HPTS, using a 1030 nm and 10 W Yb-based PHAROS laser (Light Conversion) with a pulse duration of 190 fs. It was performed at 1 MHz with a maximum pulse energy of 10 μJ. Following, an optical parametric amplifier (ORPHEUS, Light Conversion) is situated to which the laser beam will be scattered. From ORPHEUS, the laser beam is seeded into a spectrometer where the time-resolved spectrum of the sample is collected in a single photon counting technique using a hybrid detector (Becker & Hickl, HPM-100-07) that has an instrument response function (IRF) of ~50 ps at full-width-half-maximum.

**Preparation of giant unilamellar vesicles (GUVs):** POPC lipid along with $C_{12}$-HPTS (in a ratio of 100:1) were dissolved in chloroform (3 mg/ml). Following, the solution was spread on two FTO glass slides. Then the glass slides were dried using $N_2$ gas and later transferred to a vacuum desiccator for further drying. A chamber was built using dried slides and a 100 mM sucrose solution. Further, the chamber was connected to an AC generator. 1.1 V and 10 Hz sin wave were applied for the first two hours and later a sin wave of 1.1 V and 5 Hz. Finally, the solution was collected from the chamber for further analysis.

**Confocal fluorescence microscopy:** Fluorescence microscopy images were collected using an inverted confocal laser scanning microscope (LSM 710, Zeiss). The images were collected using a water objective of numerical aperture 1.2. The samples containing probe $C_{12}$-HPTS were excited using a UV laser (405 nm) and the data acquisition was performed using ZEN software.

**Table S1.** The ground and excited state pKa of all the combinations along with absorption and emission maxima for both ROH and RO$^-$ forms.

| Liposome | Absorption (nm) | | Emission (nm) | | $pK_a$ | $\Delta pK_a$ | $pK_a^*$ |
|---|---|---|---|---|---|---|---|
| | $\lambda_{ROH}$ | $\lambda_{RO-}$ | $\lambda_{ROH}$ | $\lambda_{RO-}$ | | | |
| PC | 422 | 501 | 476 | 555 | 8.07 | 7.15 | 0.9 |
| PA:PC 1:9 | 422 | 500 | 479 | 553 | 9.15 | 6.90 | 2.25 |
| PA:PC 1:3 | 423 | 500 | 479 | 552 | 9.37 | 6.81 | 2.56 |
| PA:PC 1:1 | 423 | 499 | 475 | 556 | 9.70 | 7.09 | 2.61 |
| PA:PC 3:1 | 420 | 497 | 465 | 546 | 10.03 | 7.14 | 2.89 |
| PA | 410 | 484 | 460 | 543 | 10.31 | 7.50 | 2.81 |

**Table S2.** Results from the three exponential fitting of TCSPC decay of all the mixtures and the overall average lifetime $<\tau>$.*

| | $a_1$ | $\tau_1$ | $a_2$ | $\tau_2$ | $a_3$ | $\tau_3$ | $<\tau>$ |
|---|---|---|---|---|---|---|---|
| PC | 0.63 | 0.22 | 0.22 | 1.21 | 0.15 | 3.02 | 0.85 |
| PC:PA 9:1 | 0.51 | 0.24 | 0.22 | 1.31 | 0.17 | 3.24 | 0.96 |
| PC:PA 3:1 | 0.61 | 0.23 | 0.23 | 1.26 | 0.16 | 3.12 | 0.93 |
| PC:PA 1:1 | 0.71 | 0.16 | 0.16 | 0.94 | 0.13 | 2.91 | 0.64 |
| PC:PA 1:3 | 0.75 | 0.15 | 0.17 | 0.86 | 0.08 | 3.37 | 0.52 |
| PA | 0.77 | 0.13 | 0.18 | 0.82 | 0.05 | 3.43 | 0.41 |

*All timescales are in nanoseconds.

**Table S3.** $k_{PT}$ (extracted from short time kinetics) and, B, d and $\tau_0$ (extracted from long time kinetics) for all the mixtures.

| **Liposome** | **Short time kinetics** | **Long time kinetics** | | |
|---|---|---|---|---|
| | $k_{PT}$ ($ns^{-1}$) | B | d | $\tau_0$ (ns) |
| PC | 2.86 | 2.6 | 2.7 | 4 |
| PA:PC 1:9 | 2.12 | 2.6 | 2.5 | 4.1 |
| PA:PC 1:3 | 2.65 | 2.4 | 2.5 | 3.9 |
| PA:PC 1:1 | 4.43 | 2.3 | 2.4 | 3.5 |
| PA:PC 3:1 | 5 | 2 | 1.9 | 3.2 |
| PA | 5.92 | 1.7 | 1.7 | 3 |

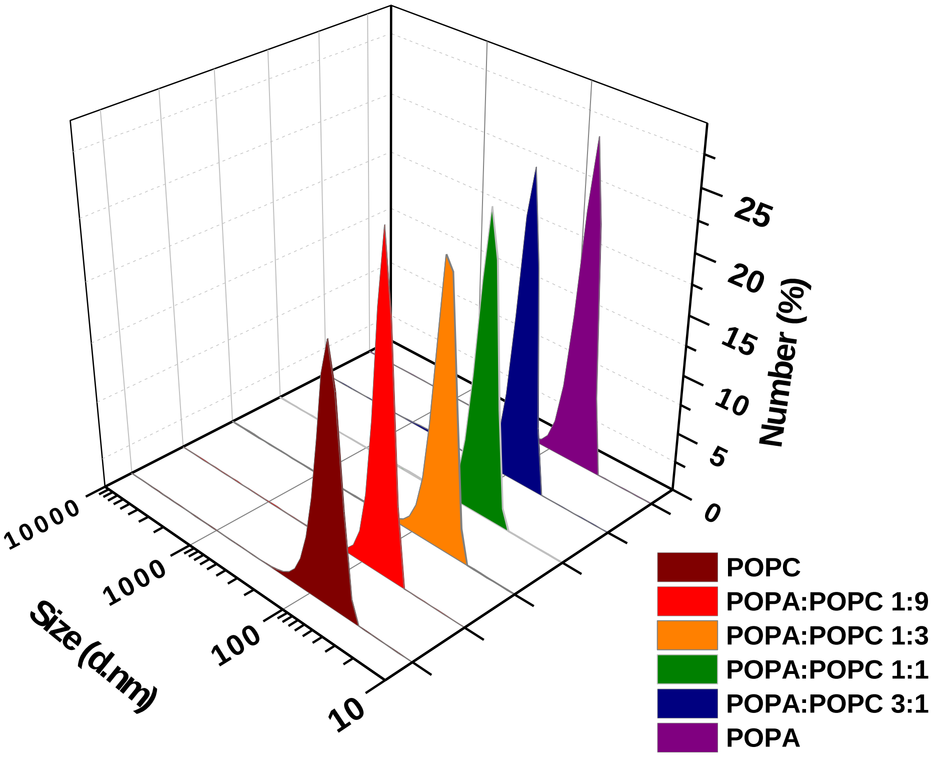


**Figure S1.** DLS shows the formation of monodispersed vesicles of size ~ 130 nm for all SUVs at different POPC:POPA ratios.

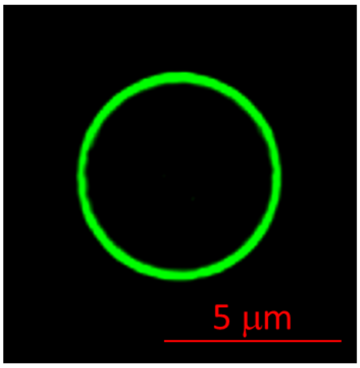


**Figure S2.** Confocal fluorescence images of POPC GUV showing the insertion of the $C_{12}$-HPTS probe into the membrane of the vesicles.

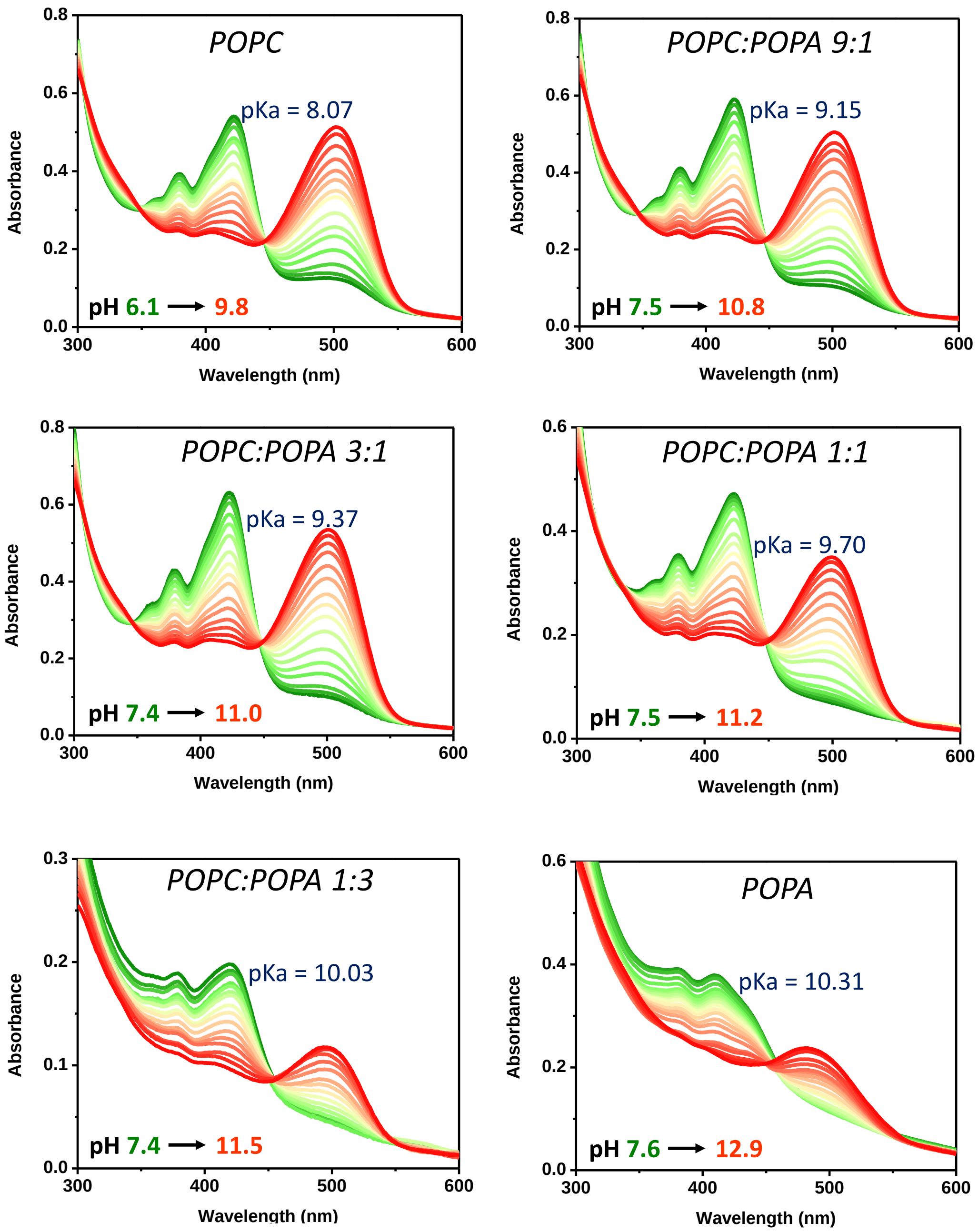


**Figure S3.** pH titration curves obtained by carrying out UV-visible absorbance experiments at different pHs for all the mixtures. The prominent peak around 420 nm and at pH ~ 7.4 represents the ROH form and the one after pH 10 and forms around 500 nm depicts the $RO^-$ form.

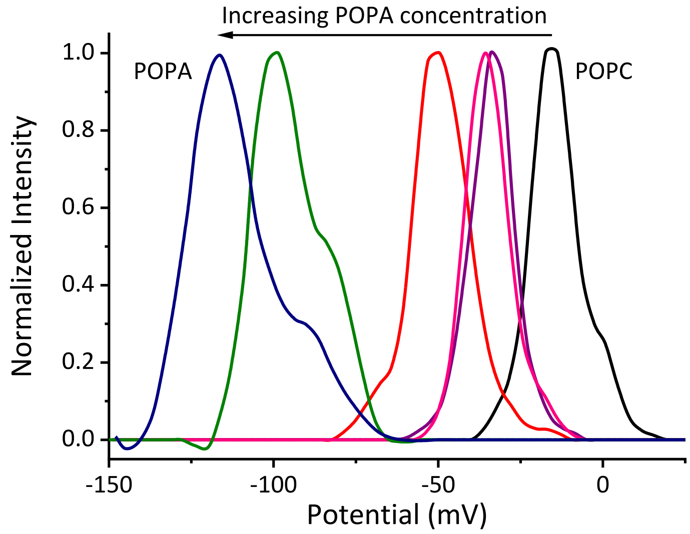

**Figure S4.** Zeta potential of all SUVs at different POPC:POPA ratios.

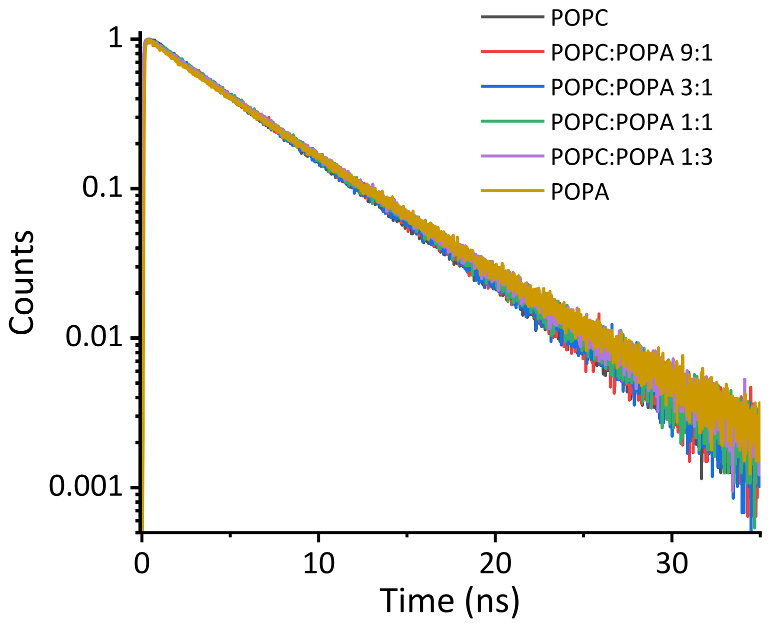

**Figure S5.** Time-resolved decay at the position of $RO^{-*}$ for all SUVs at different POPC:POPA ratios

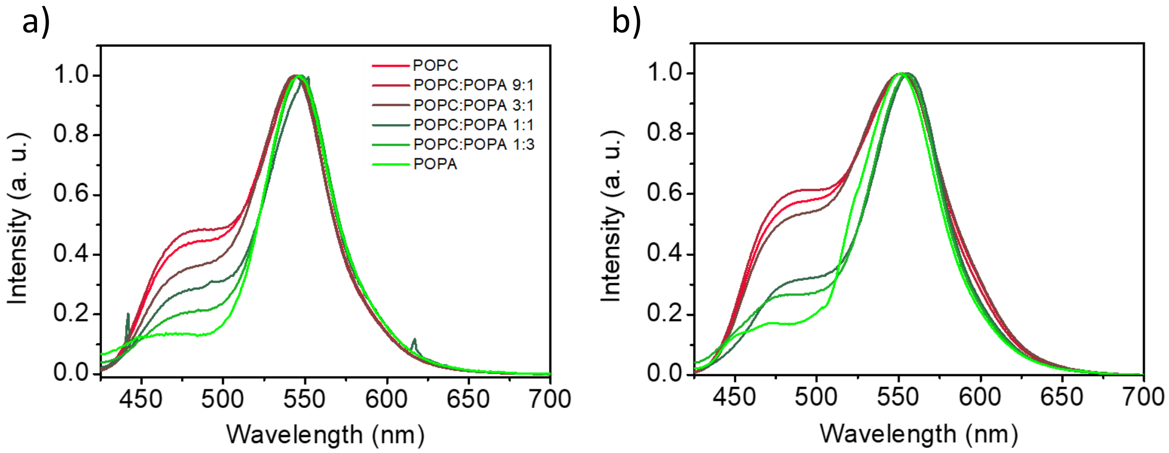


**Figure S6.** Steady-state fluorescence of all SUVs at different POPC:POPA at (a) pH 5.5 and (b) pH 3.6. All the results show the anomaly in ESPT for mixtures containing a small amount of PA.

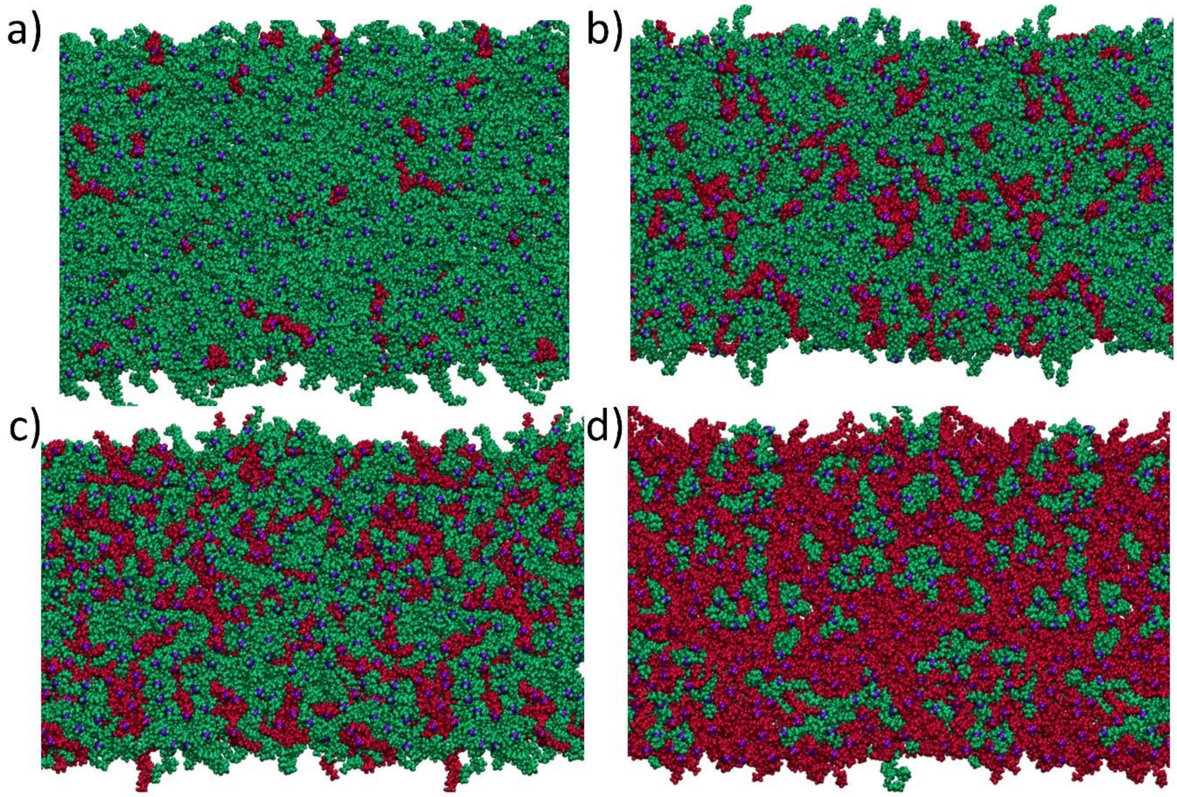


**Figure S7**. Snapshots of investigated all-atom MD membrane systems with POPC and POPA mixtures (from top view). The snapshots of (a) POPC:POPA 9:1, (b) POPC:POPA 3:1, (c) POPC:POPA 3:2 and (d) POPC:POPA 1:3 membranes are presented. Green color represents POPCs lipid and red color represents POPA lipids.

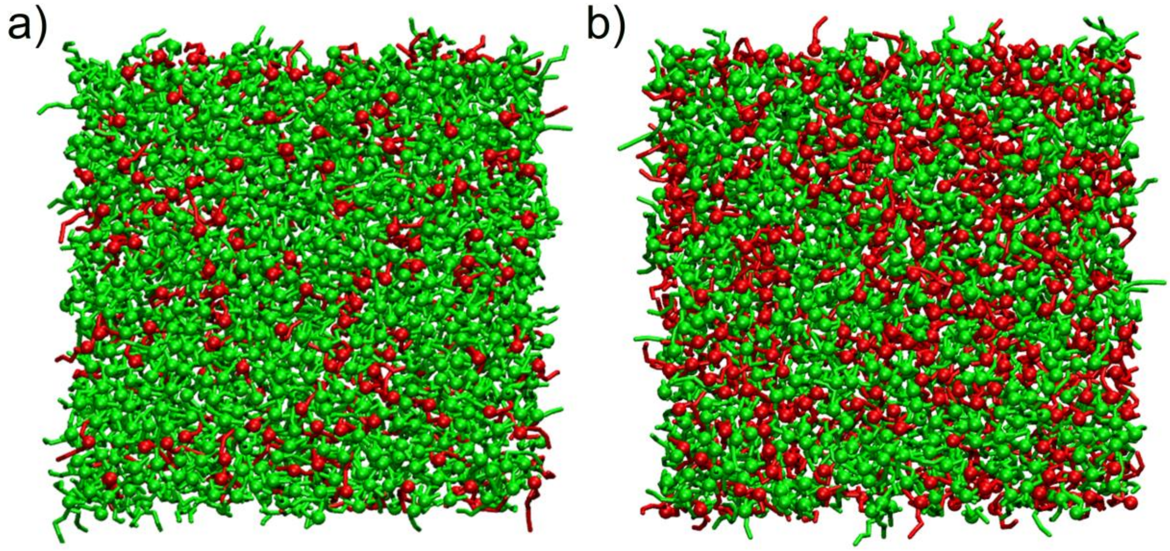


**Figure S8** Snapshots from coarse-grained membrane systems with POPC and POPA mixtures (modeled using the Martini force field (19)) from the top view. The systems comprise 1000 lipids and neutralizing ions and were equilibrated through the standard 6-step procedure from CHARMM-GUI (1). Panel A shows a system with 80% PC and 20% PA lipids, produced for 34 microseconds, while Panel B presents a system with 60% PC and 40% PA lipids, simulated for 50 microseconds. No indication of domain formation was observed in these simulations.

**Molecular Dynamics membrane systems validation:** To ensure the viability of our membrane systems, a comprehensive comparison between area per lipid (APL), membrane thickness (MT), and tail order parameter ($S_{CD}$) was performed. APL for our system ranged from (69±4) Å$^2$, (62±1) Å$^2$, (67±4) Å$^2$, (66±4) Å$^2$, (63.8±3.6) Å$^2$ and (62.0±3.3) Å$^2$ for POPC, POPC:POPA 9:1, 3:1, 3:2, 2:3 and POPA systems, respectively. These results remain consistent with existing literature. Specifically, for the POPC membrane system, an agreement was found with the literature (20-22). Similarly, for the pure POPA membrane system, the results are in line with previous research (23), although reported values are slightly lower due to simulation temperature differences. Furthermore, our outcomes notably converge with experimental monolayer studies (24, 25).

In the context of POPC:POPA mixtures, where existing literature is somewhat limited, drawing comprehensive comparisons poses a challenge. In (23) the APL for POPC:POPA 4:1 is similar to the one of the POPC system, which corresponds to our case with POPC and POPC:POPA 3:1 systems.

Similarly, MT (shown in Figure 4D) remains consistent with literature reports. Specifically, MT values of (3.7+0.2) nm and (3.95+0.15) nm were recorded for the POPC and POPA systems, respectively. The results for the POPC system fall within the range of values reported in the literature (26-28). There are limited (21) reports on membrane thickness. Thus, the values reported are slightly higher than those we report here, however, this can be attributed to variations in simulation temperatures.

Finally, values of $S_{CD}$ are reported in Figure S.9. These results remain consistent and follow the trend reported in POPC and POPA mixtures with minor deviations resulting from various force fields and temperature applications (26, 29-31)

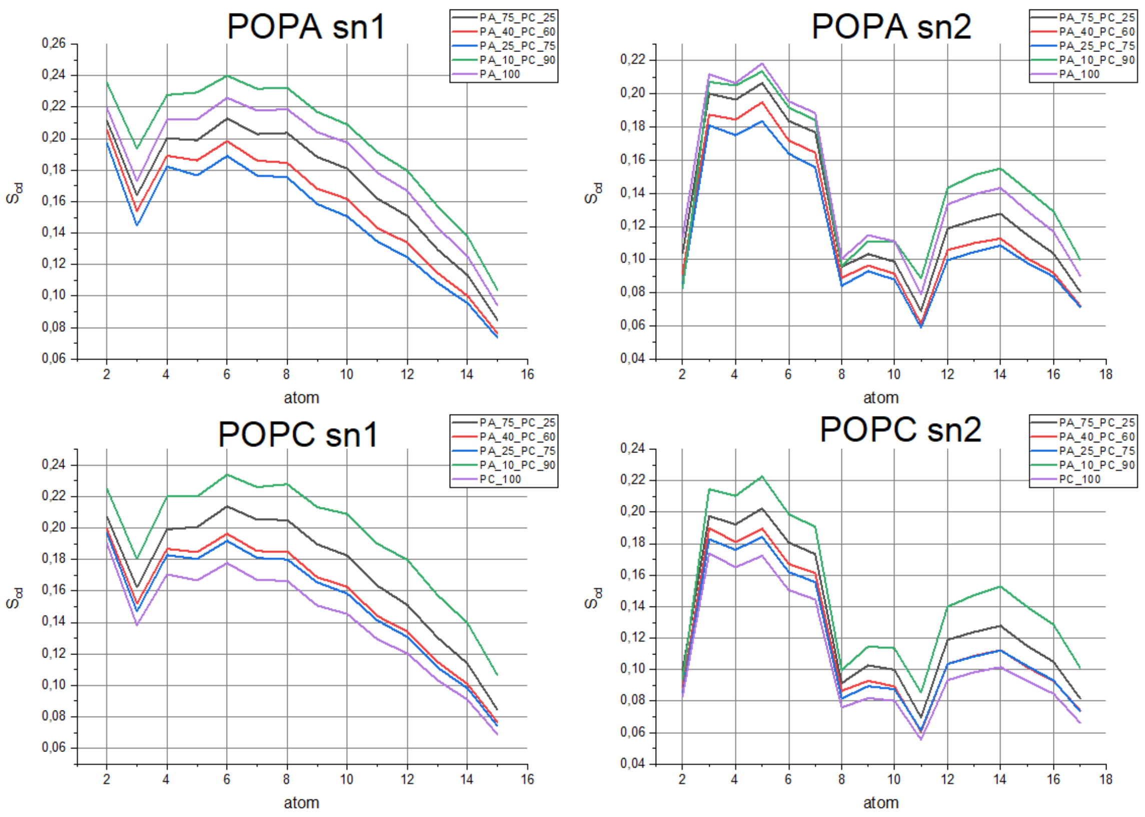


**Figure S9.** Values of $S_{CD}$ of investigated membrane systems calculated for both POPA and POPC lipid molecules for both tails (sn1 and sn2).

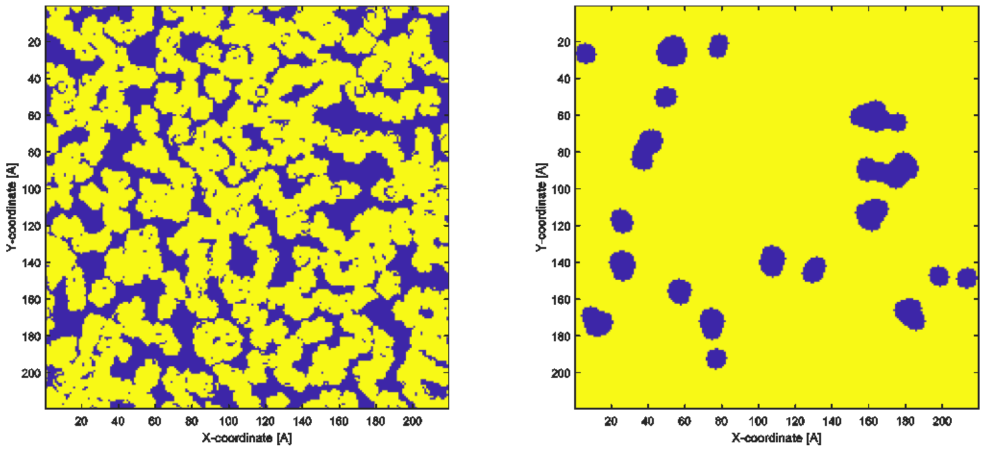


**Figure S10.** Example of defect analysis. Right panel – the grid analysis of acyl exposed regions (blue) and head molecules (yellow). Left panel – the defect analysis with sphere probe of size 2.5 Å.

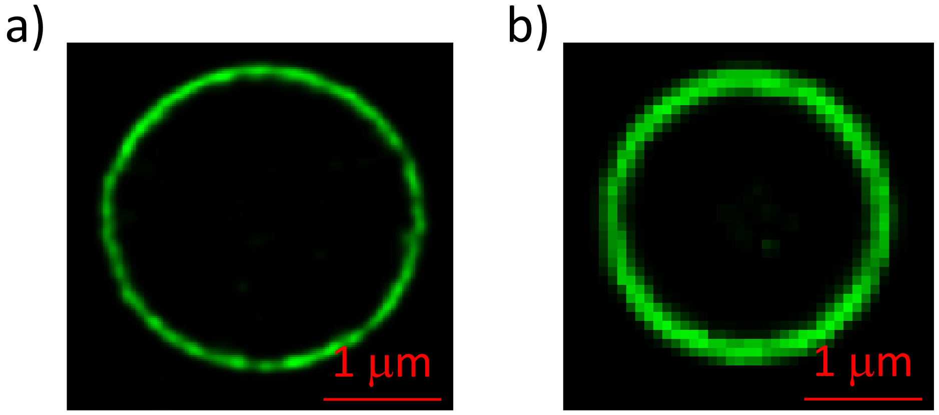


**Figure S11.** Confocal fluorescence microscopy images of (a) PC:PA 9:1 and (b) PC:PA 3:1 membrane. The pictures were taken using the same fluorescent probe $C_{12}$-HPTS.

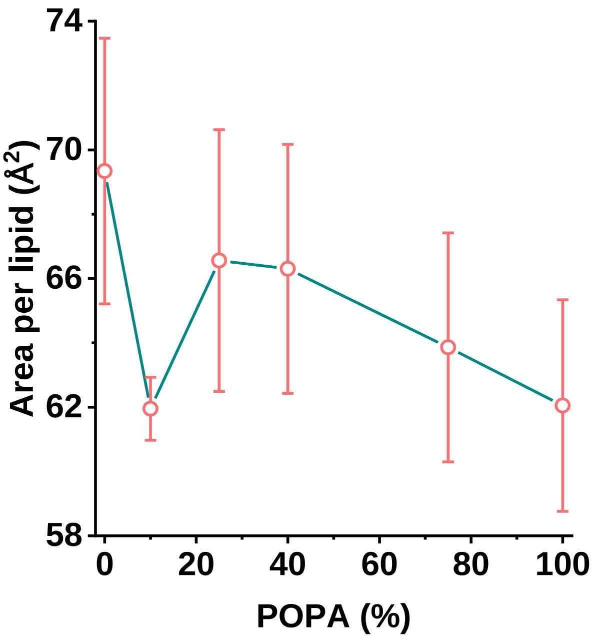


**Figure S12.** The calculated area per lipid measurements of the membranes vs % of POPA in POPC.

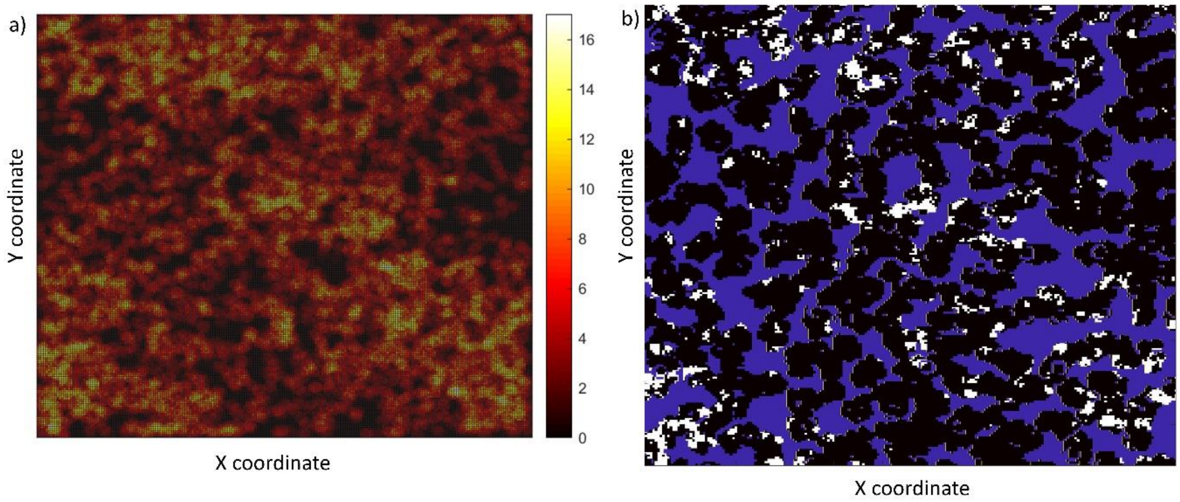


**Figure S13**. (a) Example of water slab density 1 nm above the membrane surface. (b) Image showing potential correlation of water density (white – high density, black – low density) with the position of membrane defects obtained with defect analysis. This was done for the POPC:POPA 9:1 membrane system, as this was the most structured one.

Density maps of the water slab were prepared in order to verify the proximity of higher water compaction and membrane defects. The higher the value indicated on this map, the greater the density of water molecules observed. This was followed by a threshold of the density map by half of the maximum value and superimposing the map with the defect maps. The general observation verifies our assumption, however without perfect correlation. Predominantly, defects are found in areas with a higher concentration of water molecules, yet there are some regions on the density map where this does not always hold. It is important to recognize that membranes are dynamic, curved structures, hence, this analysis of correlating water density with defect density should be taken with a grain of salt.
